# A semidwarf and late-flowering isogenic Kosihihikari d60Hd16: development, productivity, and regional suitability revealed by correlation-based network analysis

**DOI:** 10.1101/2024.06.02.597026

**Authors:** Motonori Tomita, Hiroshi Honda

**Author notes:** Corresponding author: E-mail address: Motonori Tomita.

## Abstract

Breeding rice varieties that are tolerant to weather variability and optimizing cultivation methods for each developed variety are challenging issues for global food problems. In this study, the late-flowering gene *Hd16* of Koganebare was introduced into Koshihikari through backcrossing to create ’Koshihikari Hd16’. It was then crossed with ’Koshihikari d60’ to develop an isogenic Koshihikari containing both *Hd16* and *d60*. Productivity tests were conducted in nine prefectures in Japan for two homogeneous rice genotypes, *Hd16* (late flowering) and *d60Hd16* (short culm and late flowering). By analyzing the relationship between genotype, traits, and accumulation temperature, we reexamined the characteristics of each genotype and inferred the optimal growing areas. Correlation-based network analysis among yield, grain quality, and value of taste and traits showed that quality was positively correlated with panicle length and 1000-grain weight, and yield was strongly positively correlated with 1000-grain weight. d60 genotype was negatively correlated with culm length and lodging degree. These correlations were supported by partial correlation analysis and significant differences compared to the wildtype was identified. Principal component analysis of *d60Hd16* revealed that Yamanashi and Ehime, which have longer panicle length and culm length, were suitable in terms of yield and quality, while Shimane, which is warmer and has shorter panicle length and culm length, was suitable in terms of eating quality. Moreover, Koshihikari d60Hd16 could express traits that are less prone to lodging degree while maintaining the same quality and yield as the wild type in cultivation of late-flowering strains. Thus, the *d60* and *H16* genotypes express stable traits adapted to a wide range of Japanese climatic conditions and growing environments. This study provides fundamental information for promoting new smart agriculture, in which improved varieties are deployed in different regions with different climatic conditions.

## 1. Introduction

Rice is cultivated worldwide, particularly in Asia, and is one of the world’s top three grains with an annual yield of 680 million tons, alongside corn (1.73 billion tons) and wheat (680 million tons) [1]; therefore, stable production is crucial. The earth has warmed by approximately 1.0°C from preindustrial levels, and temperatures are predicted to rise another 1.5°C between 2030 and 2052 [2]. In the IPCC’s 6th evaluation report, global warming is expected to bring about an increase in the frequency of strong tropical cyclones, and there are concerns that damage from heavy rains will be magnified [3]. Global climate change, population growth, and trade liberalization risk disrupting the current agricultural system, which could lead to food problems. Therefore, molecular genetic breeding of cereals suitable for different climatic conditions is positioned as a challenging issue to be solved.

So far, the “Green Revolution” has contributed to improving the lodging resistance of rice [4,5]. The gene for contributing rice “Green Revolution” was identified as sd1 on the long arm of chromosome 1 [6–8], encode a defective C20-oxidase in the gibberellin (GA) biosynthesis pathway (GA 20-oxidase, OsGA20ox2) [9–11] and mutations in the GA20-oxidase gene lead to disruptions at a late stage of the GA pathway [11]. The sd1 gene confers no detrimental effects on grain yield [12–14]. biol. However, the trend of rice productively has now begun to plateau [4]. Surprisingly, semidwarf rice varieties developed independently using different native varieties or artificially induced mutant lines as mother plants have the same sd1 loci. This narrow gene pool of semidwarfness has led to reduced genetic diversity of rice [13,15,16]. In preparation for future increases in population and risk of crop damage due to climate change, there is a demand for a “New Green Revolution” by renewed genetic improvements instead of *sd1*.

To identify a novel alternative gene to sd1, Tomita et al. conducted gene analyses on Hokuriku 100, a mutant line with 15 cm, and a 20% shorter culm than that of Koshihikari. Hokuriku 100 was developed through a large-scale mutation breeding by exposing Koshihikari to 20 kR of gamma radiation by ^60^Co [17]. Tomita et al. analyzed a mutation of Hokuriku 100 [18, 19] and discovered a unique heredity of a novel semidwarf gene *d60* with the gametic lethal gene gal, which complements d60, to cause gametic lethality [19]. Moreover, the isogenic line that was integrated with both d60 and sd1 derived from Jukkoku [20, 21] into Koshihikari by backcrossing [22]. The *d60sd1* line became a double recessive dwarf, indicating that *d60* is functionally independent from *sd1* and not related to the GA1 biosynthesis pathway [22]. Above all, *d60* is expected to diversify semidwarf breeding as a novel alternative of *sd1*.gene [23].

Koshihikari is additionally suffering from poor filling and widespread yield reduction caused by high temperature like heat waves. If the average daily temperature exceeds 23°C–24°C during the 20 days after heading, a white immature grain arises [24–27], both white-back immature grains and milky-white immature grains arise at 27°C, white-back immature grains occur at 30°C, and milky-white immature grains frequently happen at 33°C [28]. Recent heat waves caused 170,000 tons of high-temperature damage, namely, deterioration of rice quality, which spread and reached to 21% of the total production volume [29]. This is because the leading variety Koshihikari, which comprises 37.3% of rice acreage in Japan [30], flowers and ripens in the high-temperature phase in August. Rice industries have been strongly requiring late-flowering varieties instead of Koshihikari to avoid high temperature ripening. Tomita et al. identified late-flowering gene *Hd16* from Isehikari [31], which was identical with that from Nipponbare. If Koshihikari with both *Hd16* and *d60* genes could be developed, it would be an effective countermeasure against damage from typhoon-induced collapse and reduced production due to high temperatures.

Therefore, this study attempted to develop two lines, a late-flowering line *Hd16* and a short-stalked late-flowering line *d60Hd16*, to adapt Koshihikari to climate change. In addition, a productivity test was conducted in nine prefectures in Japan to evaluate Performance. Currently, smart agriculture using existing genotypes is being conducted to maximize yield and quality, but knowledge on the best regions and farming methods for each genotype is limited, and it is important to infer suitable cultivation areas by understanding the correlation between traits, genotypes and environmental factors. Therefore, in this study, the relationship between genotype, traits and integrated temperature was analyzed using statistical analyses such as correlation analysis, partial correlation analysis, principal component analysis and multiple comparison analysis to re-examine the characteristics, regional suitability and productivity of each genotype and to estimate the optimal growing region for each genotype. The study provides fundamental information for the development of new smart agriculture practices to deploy improved genotypes in regions with different climatic conditions.

## 2. Materials and Methods

### 2.1. Development of Koshihikari d60Hd16

Three backcrosses were conducted with Koshihikari as the recurrent parent by using a late-flowering plant compared to Koganebare that was segregated in F_2_ of Koshihikari x Koganebareas the non-recurrent parent (Figure 1). Since Koganebare is a late-flowering variety that is decent from Nipponbare, we hypothesis that the related late-flowering gene would be *Hd16*. The fourth backcrossing with Koshihikari was conducted by using the late-flowering plant that was segregated from Koshihikari*3/[Koshihikari×Koganebare F_2_ late-flowering type] BC_3_F_2_, and 90 plants of the Koshihikari*4/[Koshihikari×Koganebare F_2_ late-flowering type] BC_4_F_2_ were examined; at the same time, 60 plants of Koshihikari d60//Koshihikari*3/[Koshihikari×Koganebare F_2_] BC_4_F_2_ that were back-crossed to Koshihikari d60 were examined (Figure 1). The line Koshihikari d60 was an isogenic Koshihiakri having *d60* and *Gal*, which was developed by seven times of continuous backcrossing to a recurrent parent Koshihikari by using a non-recurrent parent of the *d60* homozygous segregant in the F_2_ of Koshihikari × Hokuriku100 [22]. A fifth backcrossing with Koshihikari was conducted with a late-flowering segregant in B_4_F_2_, which was considered *d60* homozygous; the days to heading was 64 days and culm length was 33.4 cm; 116 plants of [Koshihikari///*d60* Koshihikari//Koshihikari*3/[Koshihikari×Koganebare F_2_ late-flowering type]] BC_5_F_2_ were examined. A semidwarf segregant in BC_5_F_2_ was selected as *d60* homozygous late-flowering plant, followed by a sixth backcrossing with the Koshihikari d60, and [Koshihikari d60////Koshihikari///Koshihikari d60//Koshihikari*3/[Koshihikari×Koganebare F_2_ late-flowering type]] BC_6_F_2_ was examined. The heading date and culm length were investigated in all plants, and late-flowering and semidwarf plants that were considered *d60* homozygous were selected. Furthermore, a genetic diagnosis was conducted by the SSR marker RM16089 (chr3: 33.7 Mb), which is closely linked to the late-flowering gene *Hd16*, as we hypothesized [31].

**Figure 1.**
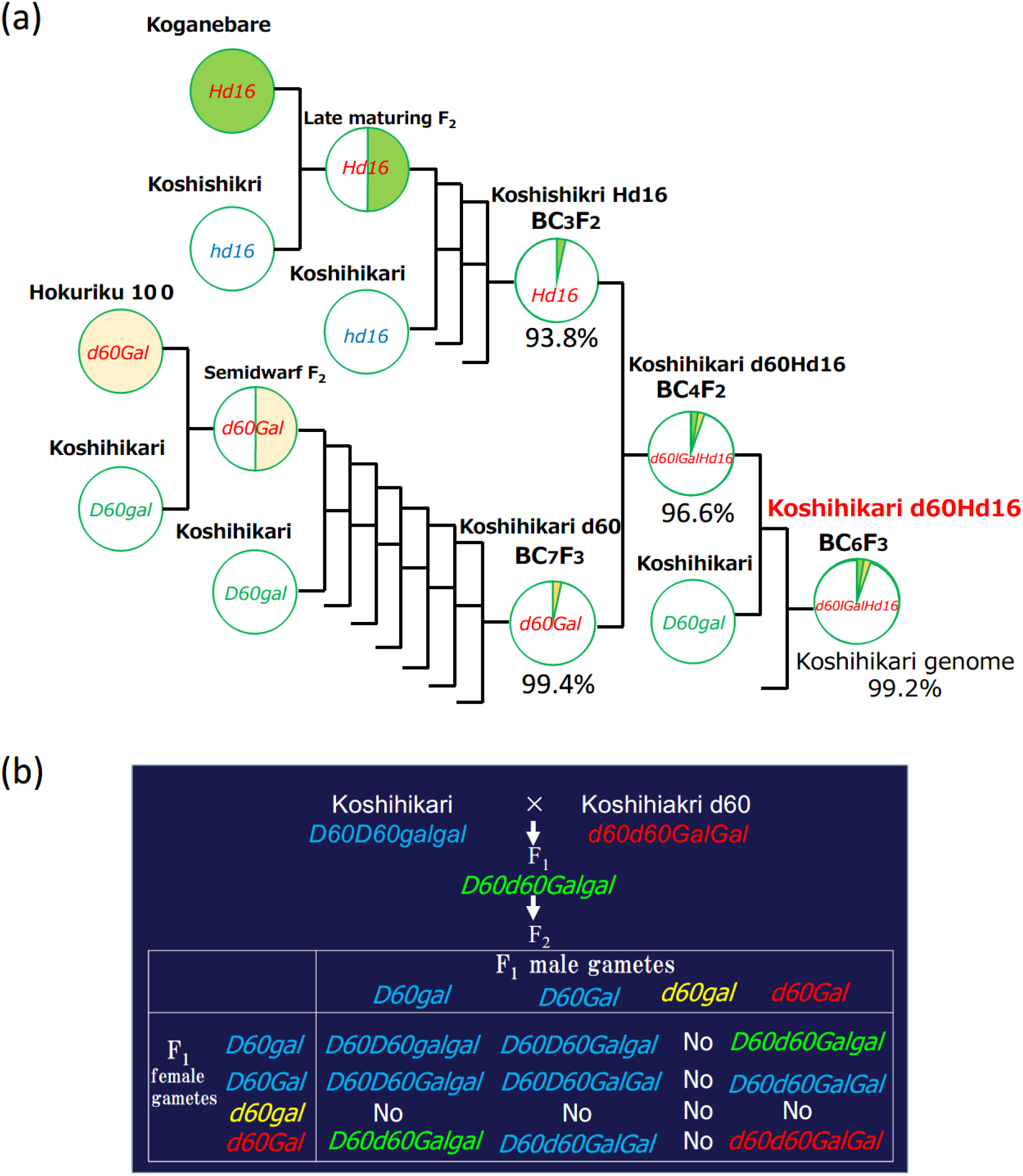
Phylogenetic process of [Koshihikari*2///Koshihikari d60//Koshihikari*3/[Koshihikari×Koganebare F_2_ late-flowering type]] BC_6_F_2_. (a) Three backcrosses were conducted with Koshihikari as a recurrent parent by using a late-flowering plant compared to Koganebare that was segregated in F_2_ of Koshihikari x Koganebare as a non-recurrent parent. The fourth backcross to Koshihikari d60 was conducted by using a late-flowering plant that was segregated from Koshihikari*3/[Koshihikari×Koganebare F_2_ late-flowering type] BC_3_F_2_. The fifth backcrossing to Koshihikari was conducted with a late-flowering segregant in B_4_F_2_, which was considered *d60* homozygous. A semidwarf and late-flowering segregant in BC_5_F_2_ selected as *d60* homozygous was sixth backcrossed to Koshihikari, and semidwarf and late-flowering BC_6_F_2_ segregants were selected as *d60* homozygous and fixed as *d60Hd16* homozygous in BC_6_F_3_. (b) From BC_4_F_2_ to BC_6_F_2_, *d60* allele was segregated in a ratio of 4*D60D60* : 4*D60d60* : 1*d60d60,* according to complementally gametic lethal with *gal* [19, 22].

### 2.2. Whole genome Sequencing Analysis

Whole genome sequencing was conducted of both Koshishikari Hd16 (BC_7_F_4_) and Koshihkari d60Hd16 (BC_8_F_2_), which was integrated with late-flowering gene *Hd16* and semidwarfing gene *sd1*, by 8 times of backcross into the genetic background of Koshihikari. The leaves were powdered using a mortar and pestle while being frozen in liquid nitrogen. The genomic DNA was then extracted from each cultivar by the CTAB method. Genomic DNA was fragmented and simultaneously tagged so that the peak size of the fragments was approximately 500 bp using the Nextera® transposome (Illumina Inc., San Diego, CA). After purification of the transposome by DNA Clean & ConcentratorTM-5 (Zymo Research, Irvine, CA), adaptor sequences, including the sequencing primers, for fixation on the flow cell were synthesized at both ends of each fragment using polymerase chain reaction, and then the DNA fragments were subjected to size selection using AMPure XP magnetic beads (Beckman Coulter, Brea, CA). Finally, qualitative checks by using Fragment Analyzer™ (Advanced Analytical Technologies, Heidelberg, Germany) and quantitative measurements by Qubit® 2.0 Fluorometer (Life Technologies; Thermo Fisher Scientific, Inc., Waltham, MA) were performed to prepare a DNA library for NGS. The sequencing was conducted in paired-end 2 × 100 bp on a HiSeq X next-gen sequencer, according to the manufacturer’s protocol (Illumina Inc., San Diego, CA). The gained Illumina reads were firstly trimmed using Trimmomatic (version 0.39) [32] (Figure 2). The sequencing adapters and sequences with low quality scores on 3′ ends (Phred score [Q], 20) were trimmed. The raw Illumina WGS reads were quality checked by performing a quality control with FastQC (version 0.11.9; Babraham Institute). Mapping of reads from Koshihikari Hd16 and Koshishikri d60Hd16 to the Koshishikri genome as a reference was conducted with Burrows-Wheeler Aligner (BWA) software (version bwa-0.7.17.tar.bz2) [33]. Duplicated reads were removed using Picard (version 2.25.5) (http://broadinstitute.github.io/picard) and secondary aligned reads removed by SAMtools (version 1.10.2) [34]. To identify genetic variations among strains, single nucleotide variant (SNV) detection (variant calling) and SNV matrix generation were performed using GATK *<* version 4.1.7.0 [35].

**Figure 2:**
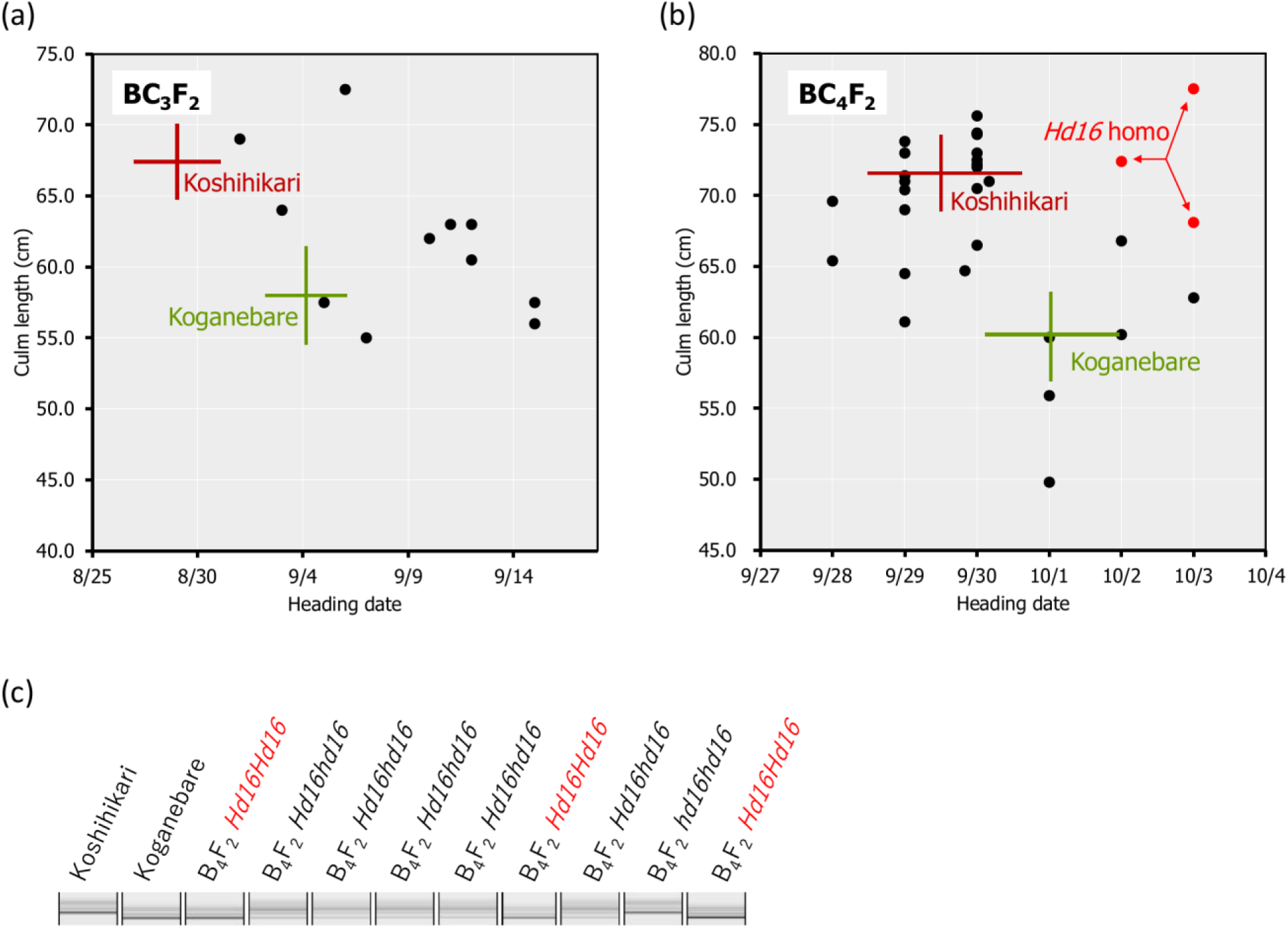
Relationship between heading date and culm length and diagnosis for *Hd16* by RM16089 in BC_3_F_2_, BC_4_F_2_ In Koshihikari*3/[Koshihikari^×^Koganebare F_2_ late-flowering type] BC_3_F_2_, which was backcrossed three times with Koshihikari as recurrent parent by using a late-flowering segregant comparing to Koganebare in F_2_ of Koshihikari ^×^ Koganebare as a non-recurrent parent, the heading dates was distributed from early-maturation comparing to Koshihikari to late-flowering later than Koganebare (a). The fourth back-crossing to Koshihikari was conducted by using the most late-flowering segregant in BC_3_F_2_. The genetic diagnosis was conducted by using the SSR marker RM16089, which is closely linked to the late-flowering gene *Hd16*, in BC_4_F_2_ of Koshihikari*4/[Koshihikari x Koganebare F_2_ *Hd16* semidwarf type](b, c). As a result, BC_4_F_2_ segregated in a ratio of 1 RM16089 homozygous: 2 heterozygous: 1 RM16089 null type and the RM16089 homozygous, namely *Hd16* homozygous isogenic line was obtained (b, c). The target genotype indicates in red. Range of culm length and heading date of each variety were shown as standard deviation using error bars.

### 2-2. productivity (performance) tests

Productivity tests were conducted in Miyagi, Yamanashi, Shizuoka, Mie, Osaka, Shimane, Ehime, Kochi and Saga prefectures for Koshihikari, Koshihikari Hd16 and Koshihikari d60Hd16 to evaluate the following parameters: culm length (㎝), panicle length (㎝), number of panicles, lodging degree, 1000 grain weight (g), protein content (%), accumulated temperature (℃), grain yield<u> (g/a)</u>, grain quality and value of taste. Accumulated temperature was treated as an environmental factor and *d60* and *Hd16* as loci. In tabulating the results of the trials, the sample name was the combination of the growing region and variety name (e.g. Yamanashi_d60Hd16). Cultivation of genetic materials was carried out in a paddy field at Shizuoka University, Shizuoka, Japan, Seedlings were individually transplanted into a paddy field in mid-July with transplanting density 22.2 seedlings/m^2^ (one seedling per 30 × 15 cm). The paddy field was fertilized by 4.0 kg of basal fertilizer containing nitrogen, phosphorus, and potassium (weight ratio, nitrogen: phosphorus: potassium = 2.6:3.2:2.6) at are with 4.3 g/m^2^ nitrogen, 5.3 g/m^2^ phosphorus, and 4.3 g/m^2^ potassium across the field. The heading date was recorded as the date first panicle had emerged from the flag leaf sheath for each plant. Culm length (㎝) was measured as the length between the ground surface and the panicle base. For the yield test, after ripening ten plants typical of each genotype were sampled twice. The sampled plants were air-dried and were assessed or measured for the following traits, panicle length, number of panicles, number of florets / panicles, proportion of fertile florets, total panicle number, and weight of not milled rice/1,000 grain. The yield of unpolished rice was calculated using the following equation: The yield of not milled rice (g/m^2^) = (number of panicles/m^2^) × (number of florets/panicle) × (proportion of fertile florets) × (weight of not milled rice/grain). Grain quality was classified into nine grade; 1: excellent good to 9: especially bad low quality. Eating taste were evaluated as seven grades of organoleptic assessment by panelist and protein contents were determined by Infratec 1241(VOSS Japan Ltd.).

### 2-3. optimal growing regions for each variety

In order to analyze the optimum growing region for each variety, the better the grain yield, grain quality and value of taste of the target traits, the more suitable the growing region was defined.

### 2-4. correlation network analysis

Correlation network analysis was conducted to visualize the correlations between the items of the productivity test results and to understand the relationships in the big picture. Pearson’s correlation coefficient and partial correlation coefficient were used as indicators of correlation in the integrated analysis of all genotypes. The partial correlation coefficient is an indicator of how two variables are correlated "without the influence of the specified variable". After first obtaining the correlation and partial correlation coefficients, the strength of the correlation coefficient between each variable was plotted as a graph of the network structure using qgraph, an R package. Relationships between variables with correlation coefficients of 0.3 or higher between variables were illustrated.

Although correlation network analysis is suitable for a broad understanding of the correlation between each variable, it is not sufficient to show the values of the correlation coefficients between variables, statistically significant differences and the certainty of the analysis. Therefore, a Pairwise scatter plot was conducted to visualize the strength of the proportional relationship between each variable when each result item of the productivity test was used as a variable. Pearson’s correlation coefficient was used to analyze the correlations, and tests of correlation coefficient and statistical significance were carried out. The analysis was carried out using the R package PerformanceAnalytics, which can simultaneously draw histograms for each variable, Pearson’s correlation coefficient and statistically significant differences between each variable and scatter plots between variables.

### 2-5. principal component analysis

Correlation network analysis and principal component analysis analyze correlations between variables, but do not clarify the relationship between each growing region and variable. Therefore, a principal component analysis was conducted to identify the relationship between each growing region (each province) and the variables that characterize it. The traits and environmental factors contributing to the separation of each sample when grouped by production area and cultivar were presented as biplots to provide insight into the optimal production area. The R package factoextra was used for the analysis.

### 2-6. multiple comparison analysis

The distribution of each trait, Environmental factors and target trait was visualized as a violin plot and the statistical significance of differences in means was analyzed using a pair wised t-test. The R package rstatix was used for the analysis.

## 3. Results

### 3.1. Development of late-maturing Koshihikari-type isogenic line “Koshihikari Hd16”

In BC_3_F_2_, which was backcrossed three times with Koshihikari as a recurrent parent by using a late-flowering segregant compared to Koganebare in F_2_ of Koshihikari x Koganebare as a non-recurrent parent (Figure 1), the heading dates was distributed from early-flowering comparing to Koshihikari to late-flowering later than Koganebare (Figure 2a). This segregation is similar to that 1 Koshihikari type early-flowering plant: 2 medium-flowering plants: 1 two weeks later flowering plants than Koshihikari caused by segregation of the *Hd16* allele from B_3_F_2_ to BC_6_F_2_ [43], in which continuously backcrossed to Koshihikari by using a late-flowering segregant in F_2_ of Koshihikari×Isehikari as a non-recurrent parent. Koganebare, which derived from the cross of Nipponbare×Kiho, is a late-flowering cultivar with the same ripening as Nipponbare. Therefore, Koganebare is presumed to harbor *Hd16* derived from Nipponbare [5]. The fourth backcrossing with Koshihikari was conducted by using the most late-flowering segregant in BC_3_F_2_. The genetic diagnosis was conducted by using the SSR marker RM16089, which is closely linked to the late-flowering gene *Hd16*, in BC_4_F_2_ of Koshihikari*4/[Koshihikari x Koganebare F_2_ *Hd16* late flowering type] (Figure 2b, c). As a result, BC_4_F_2_ segregated in a ratio of 1 RM16089 homozygous: 2 heterozygous: 1 RM16089 null type and the RM16089 homozygous, namely *Hd16* homozygous isogenic line was obtained (Figure 2b, c).

### 3.2. Development of late-flowering semidawarf Koshihikari-type isogenic line “Koshihikari d60Hd16”

Backcross with Koshishikri as a recurrent parent was conducted three times by using the Koganebare type late-flowering maturing segregant in F_2_ of Koshihikari×Koganebare as the non-recurrent parent (Figure 1). Koshihikari d60 was crossed with the latest flowering segregant in Koshihikari*3/[Koshihikari×Koganebare F_2_]BC_3_F_2_. In the BC_4_F_2_, long-culm plants (39-46 cm) similar to Koshihikari, intermediate plants, and semidwarf plants with a culm length (25-32 cm) similar to that of Koshihikari d60 and considered *d60* homozygous were segregated in a ratio of 35:5 (Figure 3a). This ratio fitted to the theoretical ratio of 8:1 caused by the segregation of *d60* and *gal* alleles. The heading date was distributed with a triadic peak in a ratio of 7 *Hd16* homozygous early-flowering type: 12 *Hd16hd16* heterozygous medium flowering type: 8 *Hd16* homozygous late-flowering type] = 1:2:1 (χ^2^ = 0.407, 0.80 <P <0.85) and this ratio was compatible with the theoretical ratio of single-gene inheritance (Figure 3b).

**Figure 3.**
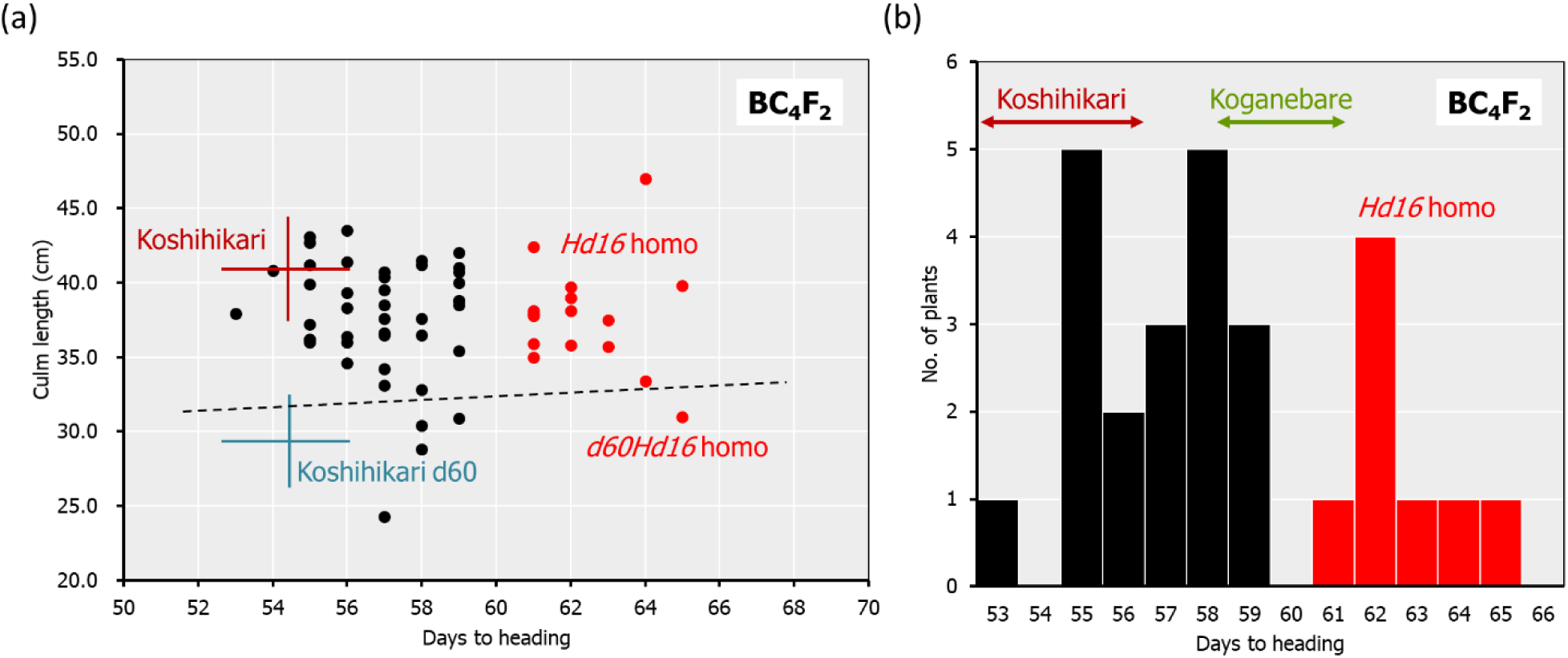
Relationship between heading date and culm length in BC_4_F_2_ of Koshihikari d60//Koshihikari*3/[Koshihikari^×^Koganebare F_2_] In BC_4_F_2_, long-culm plants (39-46 cm) similar to Koshihikari, intermediate plants, and semi-dwarf plants with a culm length (25-32 cm) similar to that of Koshihikari d60 and considered to be *d60* homozygous were segregated in a ratio of 35:5 (a). The segregants under hash line indicates *d60* homozygous plants. This ratio fitted to the theoretical ratio of 8:1 caused by the segregation of *d60* and *gal* alleles. The heading date was distributed with triadic peak in a ratio of 7 *hd16* homozygous early-flowering type: 12 *Hd16hd16* heterozygous medium flowering type: 8 *Hd1*6 homozygous late-flowering type (red)] = 1:2:1 (χ^2^ = 0.407, 0.80 <P <0.85) and this ratio was compatible with the theoretical ratio of single-gene inheritance (a, b). The *Hd16* homozygous plants indicates in red. Range of culm length and days to heading of each variety were shown as standard deviation using error bars.

The *d60* homozygous late-flowering segregant in BC_4_F_2_ with days to heading 64 days and culm length 33.4 cm was fifth backcrossed with Koshihikari. In the BC_5_F_2_, long-culm plants (58-71 cm) like Koshihikari, and semi-dwarf plants (∼54 cm) like the Koshihikari d60 were segregated in a ratio of 21:1 and this ratio was fitted to the theoretical ratio of 8:1 caused by *d60* and *gal* alleles. Genetic diagnosis of *Hd16* was conducted by using the SSR marker RM16089, and a *d60Hd16* homozygous semidwarf plant (54 cm, 10/7) was selected, and furthermore it was sixth backcrossed to Koshihikari. In the BC_6_F_2_, long-culm plants (56-65 cm) comparing to Koshihikari or intermediate plants, and semidwarf plants (44∼49 cm) comparing to the Koshihikari d60 were segregated in a ratio of 26:5 and this ratio was according to the theoretical ratio of 8:1 (Figure 4a). Genetic diagnosis of *Hd16* was conducted in BC_6_F_2_ and BC_6_F_3,_ by using the SSR marker RM16089, and a *d60Hd16* homozygous semidwarf BC_6_F_2_ plant (46 cm, 10/14) was selected (Figure 4a, b), and all progenies were fixed with *d60* homozygous semidwarf type in BC_6_F_3_ (Figure 4b, c). As a result, Koshihikari d60Hd16 was obtained. The semidwarf late-flowering Koshihikari isogenic line having *d60* and *Hd16* homozygous acquired in BC_6_F_3_ was a 14.6 cm (16%) shorter than Koshihikari and was characteristic with deep green color (Fig. 4d).

**Figure 4.**
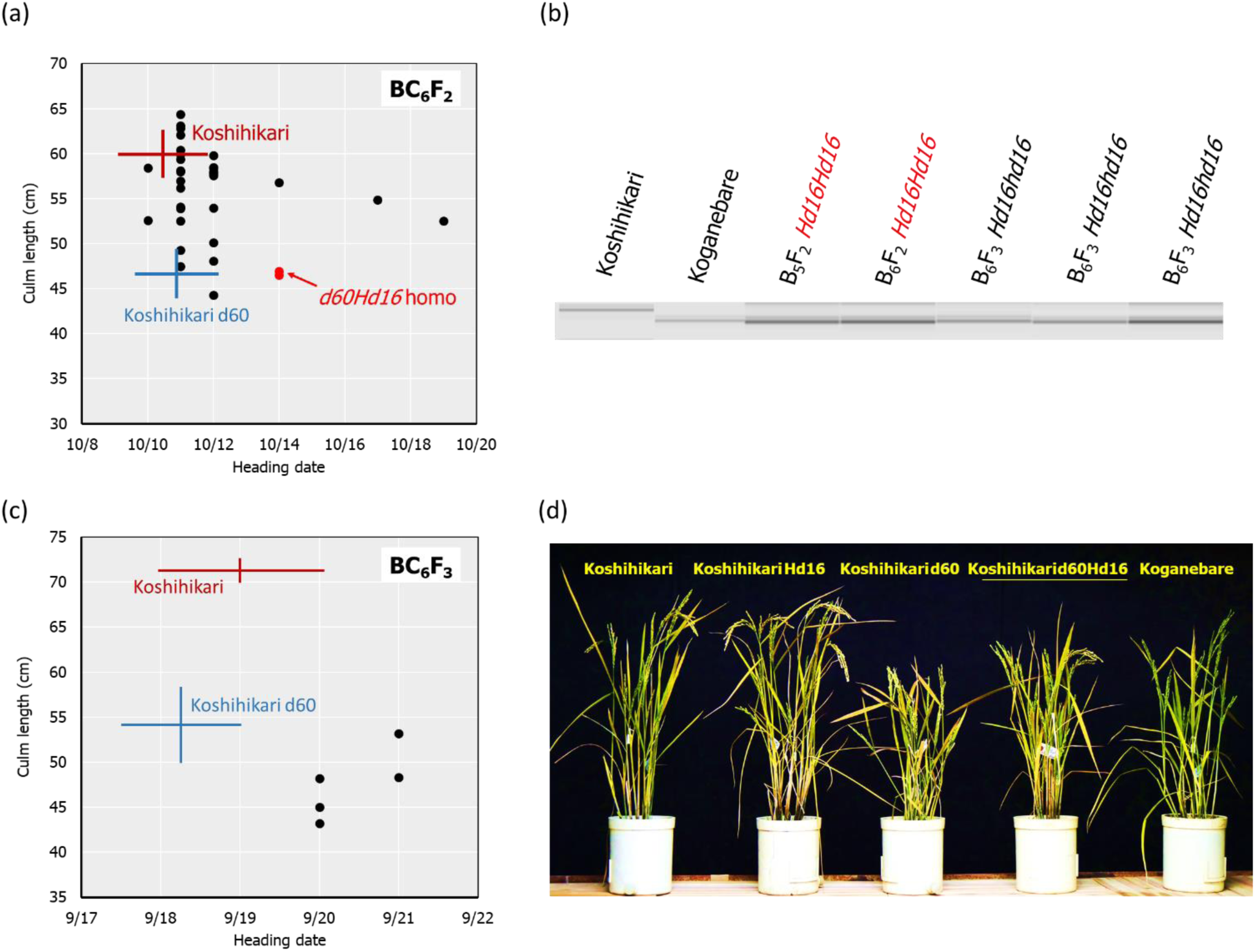
Relationship between heading date and culm length, diagnosis and morphological alteration in BC_6_F_2_ and BC_6_F_3_ In BC_6_F_2_, long-culm plants (56-65 cm) comparing to Koshihikari or intermediate plants, and semi-dwarf plants (44∼49 cm) compared to Koshihikari d60 were segregated in a ratio of 26:5 and this ratio was fitted to the theoretical ratio of 8:1 (a). Genetic diagnosis for *Hd16* was conducted by using the SSR marker RM16089, and a *d60Hd16* homozygous semi-dwarf BC_6_F_2_ plant (46 cm, 10/14) was selected (a, b). The target genotype indicates in red. The progeny line BC_6_F_3_ was fixed in *d60Hd16* homozygous (b, c). The semidwarf late-flowering Koshihikari isogenic line having *d60* and *Hd16* homozygous acquired in BC_6_F_3_ was a 14.6 cm (16%) shorter than Koshihikari and was characteristic with deep green color (d). Range of culm length and heading date of each variety was shown as standard deviation using error bars.

In Koshihikari d60Hd16 (BC_6_F_3_), SNPs from adenine to guanine, were detected in *Hd16* gene at 32,996,608 bp on chromosome 3, which was known as a causative mutation of *Hd16* in Nipponbare [5] and Isehikari [43] (Figure 5). Except for the region around *Hd16*, the number of SNPs were less than 10 per 0.1 Mb (Figure 5). The results indicated that a large portion of the rice 12 chromosomes were substituted to the genome of Koshihikari after continuous backcross targeting *Hd16*.

**Figure 5.**
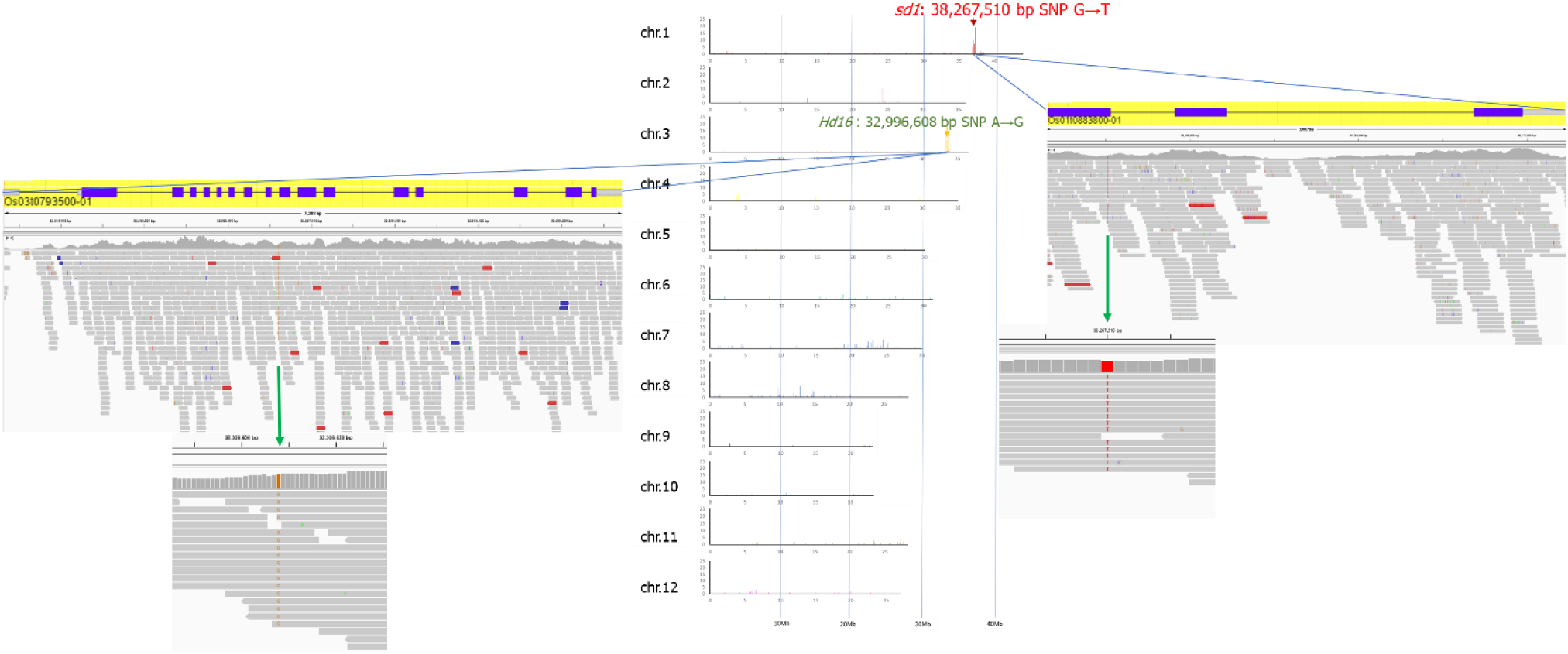
Causative SNP for *Hd16* in Koshihikari d60Hd16. A single SNP from adenine to guanine, was detected at 32,996,608 bp in *Hd16* gene on chromosome 3, which was known as a causative mutation of *Hd16* in Nipponbare.

### 3.3. Correlation and partial correlation analysis

The results of the productivity test trials integrating the three varieties are shown in Supplementary table S1. The data of each variety were integrated and the relationship between yield, grain quality, value of taste and traits were analyzed. temperature (Supplementary table S1, Figure 6). A positive correlation was also found between grain quality and panicle length in the analysis of partial correlation coefficients. Grain yield was strongly positively correlated with 1000 grain weight in the correlation analysis. These correlations were confirmed in the partial correlation analysis, which also found positive correlations with lodging degree and culm length, making it easy to understand that the heavier the grain yield, the easier the lodging degree. The correlation analysis did not show a significant correlation for value of taste, but the partial correlation analysis showed a negative correlation with culm length and protein content. Negative correlations were found with culm length and protein content in the partial correlation analysis.

**Figure 6.**
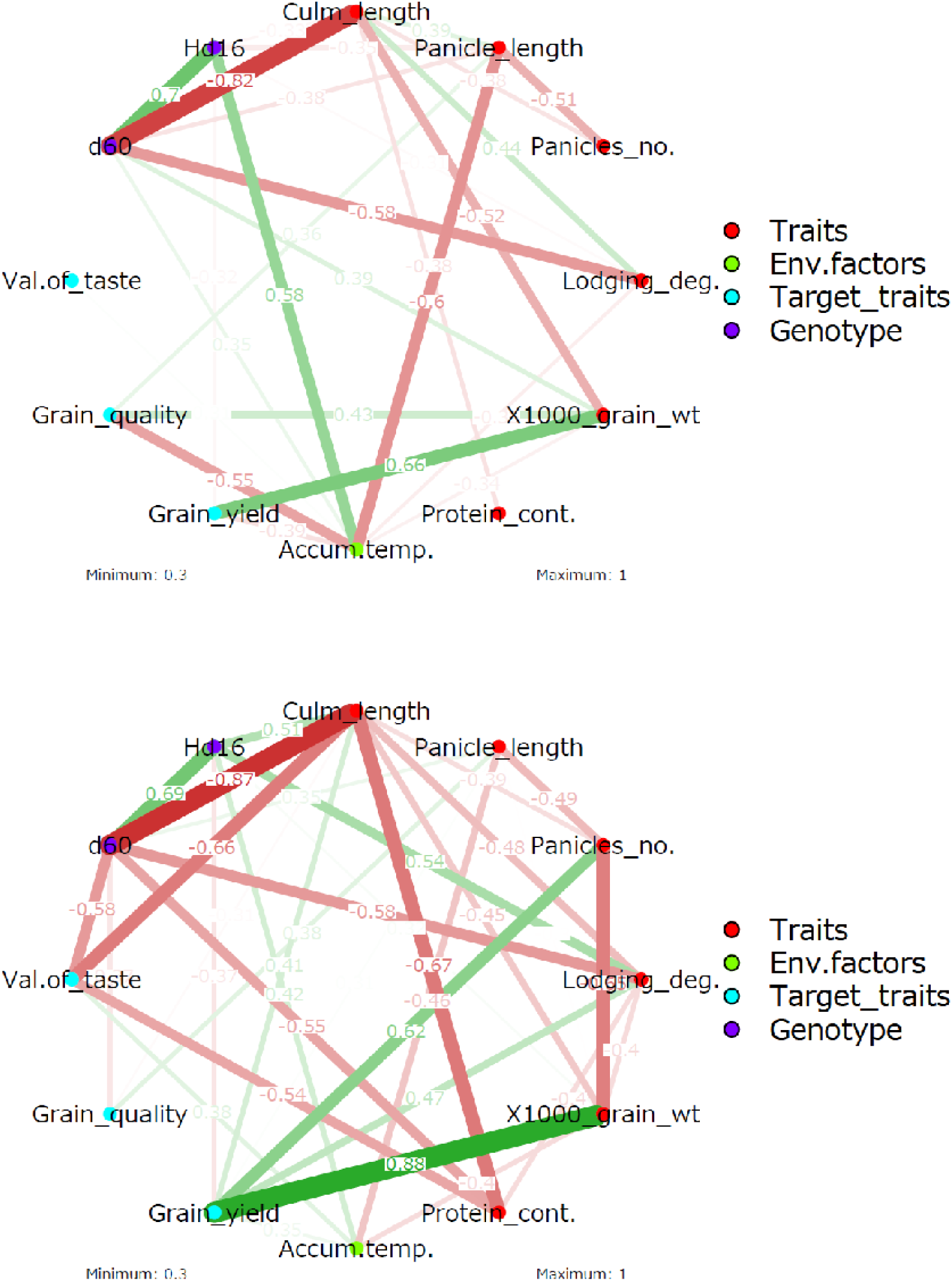
Correlation-based network analysis on traits, environmental factors, target traits, and genotype. A) correlation analysis, B) partial-correlation analysis. WT: wildtype, Panicles_no.: No. of panicles, Lodging_deg.: Lodging degree, 1000_grain_wt: 1000 grain weight, Protein_cont.: Protein content, Accum.temp.: Accumulated temperature. The values on edges indicates Pearson correlation coefficient. The density of the network indicates the strength of the correlation coefficient, with red indicating a negative correlation and green indicating a positive correlation.

Next, the relationship between genotype and trait was confirmed. In correlation analysis, *Hd16* was negatively correlated with panicle length. However, *Hd16* was positively correlated with accumulated temperature due to the difference in cultivation time, and the negative correlation between accumulated temperature and panicle length indicated that the negative correlation between *Hd16* and panicle length was a spurious correlation. No trend was observed in the partial correlation analysis. In the partial correlation analysis, correlations between traits should be considered, but positive correlations with culm length and lodging degree and negative correlations with grain yield were found. In correlation analysis, *d60* was found to be negatively correlated with culm length and lodging degree. Furthermore, in partial correlation analysis, *d60* was negatively correlated with culm length and lodging degree, and newly negatively correlated with value of taste and protein content. This negative partial correlation with value of taste and protein content is interpreted as originating from the negative partial correlation between culm length and value of taste and protein content.

The scatter plots between variables and the statistical significance of the correlation coefficients are shown in Supplementary figure S1. It was confirmed that the correlation coefficients were statistically significant at the high correlation coefficient locations and that the distribution of the samples was uniform in the scatter plots, thus confirming that there were no problems with the reliability of the correlation analysis.

### 3.4. Principal component analysis

The results of the principal component analysis of the productivity test trials integrating the three varieties showed that they were arranged separately according to genotype based on the principal component 1 and 2 (Figure 7a). The wild-type genotypes tended to have longer complement lengths and were more prone to lodging degree than the other genotypes, but appeared to have more grain quality, protein content, 1000 grain weight, grain yield, etc. Acquisition of the *Hd16* genotype tended to increase culm length, decrease protein content and improve value of taste, but there was greater variability and the number of panicles, 1000 grain weight and grain yield tended to be lower. This is probably due to the late maturing traits. This trend was probably influenced by the change of growing season to one more suited to the late trait. On the other hand, the acquisition of *d60* by this Koshihikari Hd16 resulted in a reduction in variability and a recovery of grain quality, 1000 grain weight and grain yield to the same levels as wild-type Koshihikari. It was verified that even when cultivated for late maturity, grain quality was maintained at the same level as wild-type Koshihikari, and the trait of resistance to lodging degree was acquired by shortening the culm length.

**Figure 7.**
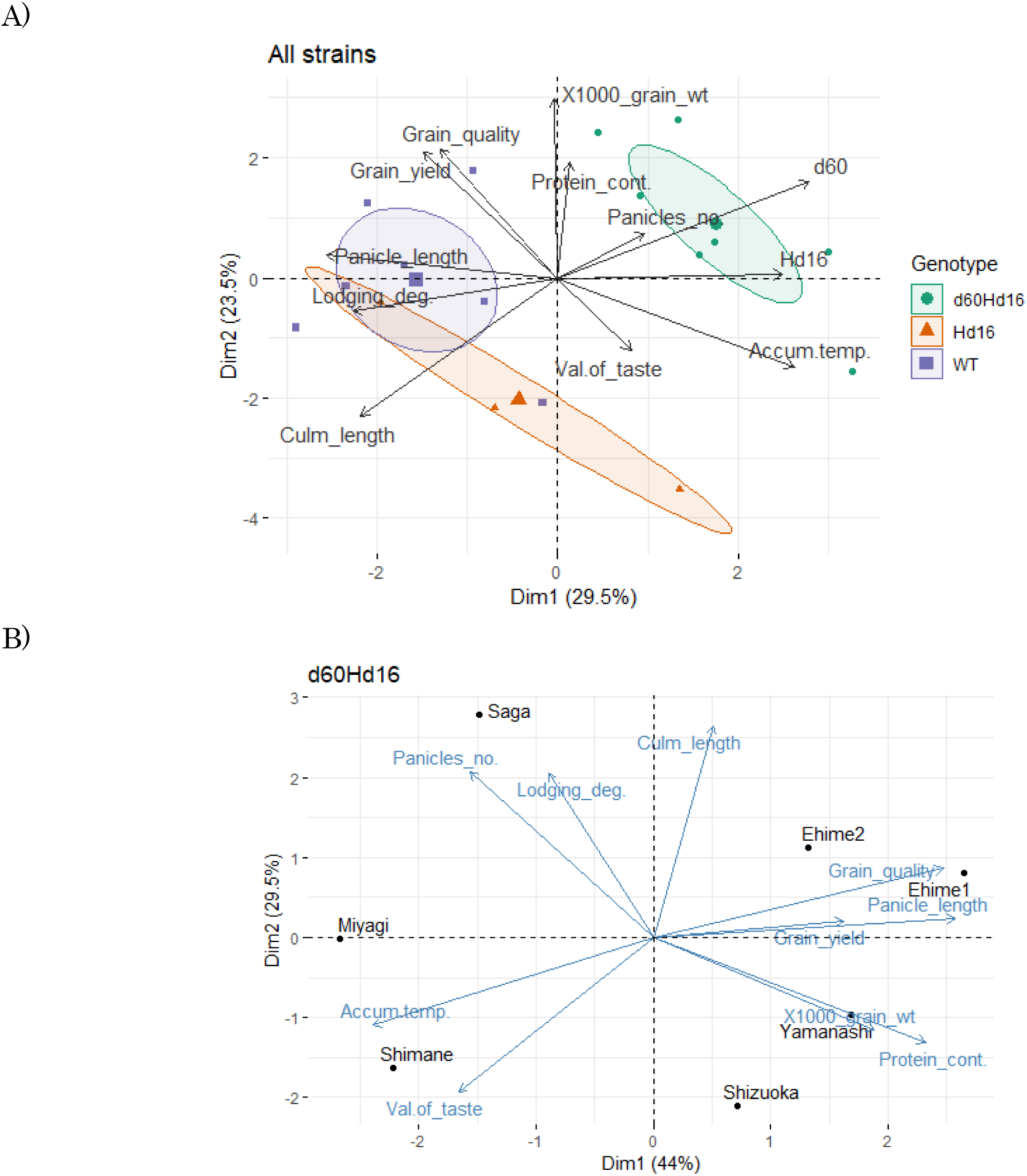
Principal component analysis for analyzing features differences among genotype. A) three strain data, B) *d60Hd16* data. Dim1: dimension 1 (Principal component 1), Dim2: dimension 2 (Principal component 2). WT: wildtype, Panicles_no.: No. of panicles, Lodging_deg.: Lodging degree, 1000_grain_wt: 1000_grain_weight, Protein_cont.: Protein content, Accum.temp.: Accumulated temperature.

Next, analysis of cultivation areas suitable for *d60Hd16* showed that Yamanashi and Ehime, which have longer panicles and culm lengths, are better from the viewpoint of yield and grain quality, while Shimane, which is warmer and has shorter panicles and culm lengths, is better from the viewpoint of value of taste. Shimane, which is warmer and has shorter panicle length and culm length, was found to be better from the viewpoint of value of taste (Figure 7b).

### 3.5. Multiple comparison analysis

The distribution of each trait, environmental factors and target trait was visualized in a violin plot and statistically significant differences in mean differences were analyzed (Supplementary figure S2 and Figure 8). To further confirm the short culm length and low lodging degree of *d60Hd16* shown in the correlation and principal component analyses, the differences in trait values between genotypes were checked. The results showed that the culm length was significantly longer in *Hd16* and significantly shorter in *d60Hd16* compared to the wild type. It thereby indicated that a lodging degree was less likely to occur in *d60Hd16* compared to the wild type, although correction for multiplicity eliminated the significant difference. No other statistically significant differences in the target traits were found, suggesting that they are equivalent to the wild type. These results suggest that the Koshihikari d60Hd16 trait can be stably expressed in a wide range of regions in Japan.

**Figure 8.**
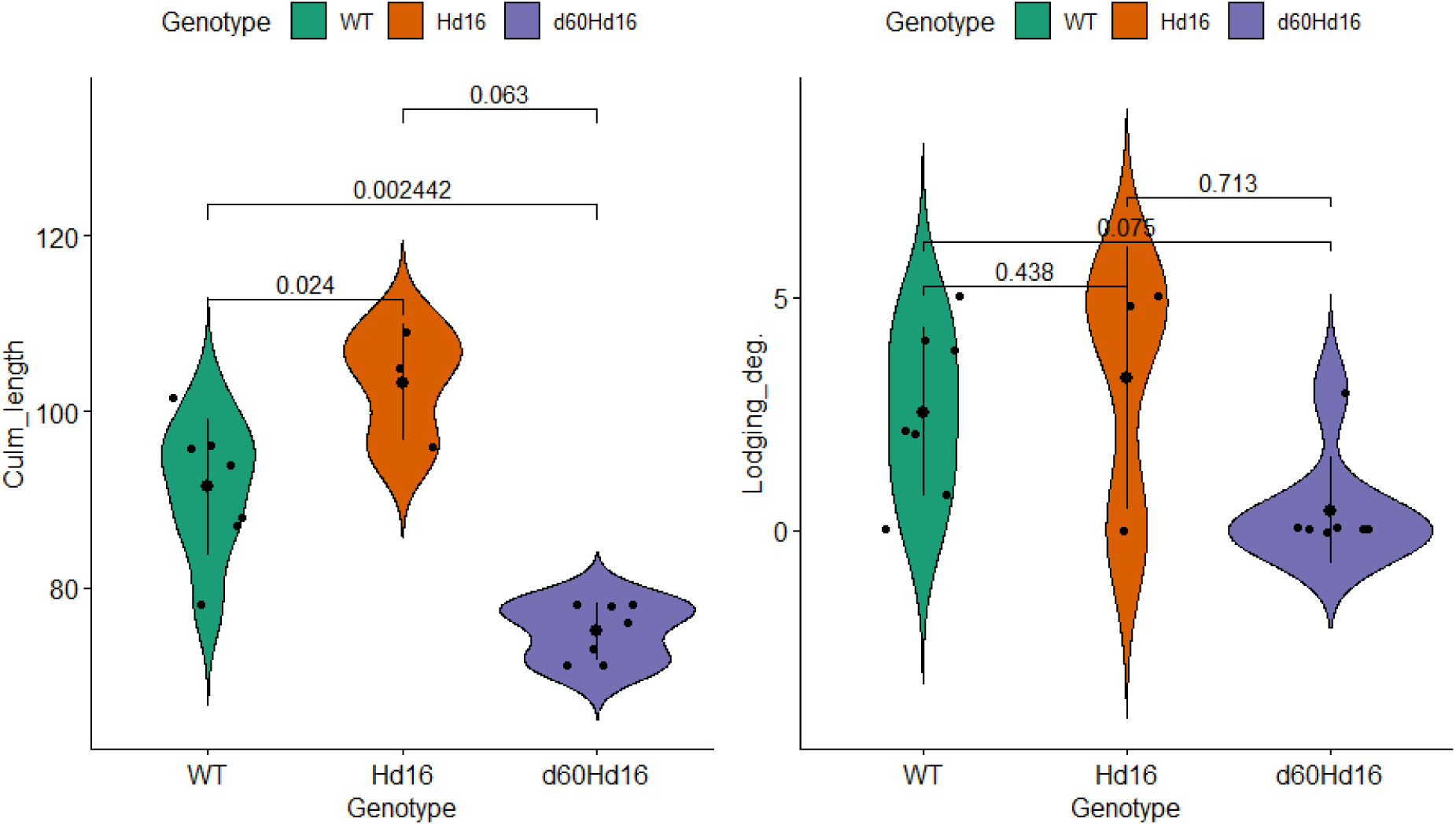
comparison of traits among genotypes. Small black points indicate individual data. Large points and bar indicate mean ± SD. Adjusted p-values using holm method are shown on the horizontal bar. Lodging_deg.: lodging degree. Lodging degree was determined based on the inclination angle of plant; 0: standing, 1: almost 70, 2: almost 50, 3: almost 30, 4: almost 10, 5: lodged.

## 4. Discussion

The present study indicates four major conclusions; 1) rice two isogenic genotypes, *Hd16* (late flowering) and *d60Hd16* (short culm and late flowering) were built in homogeneous background of Koshishikri to adapt Koshihikari to climate changes. 2) quality was positively correlated with panicle length and 1000-grain weight, and yield was strongly positively correlated with 1000-grain weight in correlation and partial-correlation analysis, 3) The *d60* genotype was negatively correlated with culm length and lodging degree, and has the effect of improving the risk of collapse (increase of culm length) and low yield that might be caused by *Hd16* genotype, and 4) suitable cultivating regions were suggested for each variety in PCA.

The threat of strong typhoons, rainfall, and flood caused by global warming is [36] causing serious lodging, consequent yield loss and grain quality deterioration in rice production [37]. A novel dwarf gene, *d60*, which was found in the semidwarf mutant Hokuriku 100 [18, 38], is thus of particular importance. *d60* is inherited as a complemental gene with a gametic lethal gene gal. Consequently, the cross between Hokuriku 100 and Koshihikari exhibits a unique genotype ratio of 4*D60D60* : 4*D60d60* : 1*d60d60* [19, 22]. In addition, Koshihikari is also suffering from poor filling and yield reduction caused by high temperature maturation. To avoid high-temperature damage in the hot summer, it is an effective solution to shift rice ripening to early autumn. In this study, firstly, the late-flowering gene *Hd16* from Koganebare was integrated to Koshishikri by four times of backcrossing with Koshihikari as a recurrent parent by using a late-flowering plant as a non-recurrent parent that was segregated in F_2_ of Koshihikari x Koganebare. Furthermore, in this study we developed a semidwarf late-flowering Koshihikari-type isogenic line (Koshihikari d60Hd16), for the purpose of stabilizing high yield and to avoid high temperature maturation. Namely, the late flowering isogenic Koshishikri Hd16 was crossed with Koshihikri d60 to combine semidwarf gene *d60* and *Hd16* into the genetic background of Koshihikari, and finally six times of backcrossing to the genetic background of Koshihikari was completed to build isogenic Koshihikari integrating both *Hd16* and *d60*. Through the process, *Hd16* allele was diagnosed by SSR marker RM16089 near *Hd16* allele and *d60* allele was selected by its unique phenotypic segregation ratio according to 4*D60D60* : 4*D60d60* : 1*d60d60* by testing relativity limited each BCnF2 population. Finally, whole genome sequencing of Koshishikri d60Hd16 proved that a SNP from adenine to guanine was detected at 32,996,608 bp in *Hd16* gene on chromosome 3, which was the same as a causative mutation of *Hd16* in Nipponbare. Through the process, *Hd16* allele was diagnosed by SSR marker RM16089 and *d60* allele was phenotypically selected by its unique segregation according to 4*D60D60* : 4*D60d60* : 1*d60d60*. Finally, whole genome sequencing of ‘Koshihikari d60Hd16 proved that almost genome was substituted to Koshihikari genome except for *Hd16* region, in which a causative SNP of *Hd16* at 32,996,608 bp on chromosome 3. Koshihikari d60Hd16 was 14.6 cm (16%) shorter and 12.4 days late flowering than Koshihikari. The yield of Koshishikari d60Hd16 (62.6 kg/a) was 6.0% higher than that of Koshihikari. Koshihikari d60Hd16 was registered as “Koshishikri Suruga d60Hd16” [39].

Data from productivity tests has been used for understanding the relationships between traits, qualities, cropping regions, and environmental factors [40–52]. For instance, certain rice varieties show superior grain appearance under high-temperature conditions [42] (Ishimaru et al.). AI-Daej (2022) [40] reported correlations among quality and mineral element contents. Xu et al. (2015) [52] showed that 1000-grain weight negatively affected quality but positively correlated with yield. In the present study, 1000-grain weight was positively correlated with both quality and yield, but partial correlation analysis suggested that the correlation with quality was indirect (no partial correlation was found). Li et al. (2019) [45] reported that the filled grain number per panicle, plant height, panicle length, grains per panicle, long growth period, low panicle number have accounted for high yield. Plant height had a positive effect on yield for the Indica and Japonica inbred, but had a negative effect for the japonica hybrid. In the present study, panicle length was not associated with the yield, culm length and panicle number were positively related with yield (only in partial-correlation). Correlations observed between multiple traits may only detect indirect correlations; however, to our knowledge, there are no reported cases of relationships between traits being tested by partial correlation. Thus, the present study stringently confirmed the correlation between the traits and environmental factors and provided fundamental knowledge to plan efficient cropping. The 1000-grain weight is an important trait for estimating high yield, and number of paddy and culm length are considered to be meaningful traits in terms of partial correlation.

The present study demonstrated that the varieties evaluated were suggested to express stable traits in a wide range of regions across the country. Curiously, although sample size is very small, *Hd16* was related to not only late flowering, but also increase of value of taste and culm length. This novel observation was explained by the correlation between accumulated temperature with value of taste, and partial-correlation between that and culm length. By using partial-correlation, it was suggested that accumulated temperature could affect culm length. It is probably a secondary effect of the change in cropping season and *Hd16* itself would not have the ability to control such traits. On the other hand, *d60* has the effect of improving the risk of collapse (increase of culm length) and low yield that might be caused by *Hd16* genotype. The fact that the acquisition of *d60* maintains the late flowering traits of *Hd16*, and improves large inter-regional variation and low yields, is an interesting finding and suggests that the *d60* and *Hd16* combination is extremely meaningful for the development of new varieties for adapting to climate change.

The present study clearly showed the difference of genotypes and cultivating regions using PCA. Similar analysis using PCA to analyze relationships among different strains of resistant isogenic lines in Koshihikari was conducted by Ishizaki et al. (2005) [43] and those isogenic lines have similar agronomic traits, quality, and taste with the original variety. Likewise, Semeskandi et al. (2024) [48] showed the usefulness of combination analysis using PCA and correlation analysis for rice improvement cultivation. Whereas, to our knowledge, few studies were found to use PCA to identify differences among regions cultivated. Our results suggested suitable cultivating regions for each strain and found that Yamanashi and Ehime, which have longer panicles and culm lengths, are suitable in terms of yield and grain quality, while Shimane, which is warmer and has shorter panicles and culm length is preferable for value of taste. This result is understandable by the results of correlation analysis in the present study. Value of taste was negatively correlated with grain quality and accumulated temperature was positively correlated with the quality, but was negatively correlated with the quality and yield. Thus, in warmer regions, the taste value is higher, while quality and yield are lower.

The present study does not clarify the difference of traits in regions with similar accumulated temperature. It has been investigated relationship between environmental factors and traits and some studies highlight the importance of environmental conditions in determining yield and quality. Genotypic and phenotypic correlation are noted (Tiwari et al.) [51], as well as the impact of high temperatures on grain appearance (Ishimaru et al.) [42] and the effect of water consumption and rainfall on yield (Semeskandi et al.) [48]. Therefore, adding various environmental factors, such as precipitation, to the new analysis items may provide a deeper understanding of regional factors influencing traits.

If a link between genotype and trait, as well as various environmental factors, is found, it is expected that quantitative simulation of yield in new growing areas, e.g. by modeling the relationship, can be carried out. For the prediction of traits, it is important to expand the spatio-temporal data, it is necessary to obtain data on different climatic conditions at the same location, as well as regional characteristics. Therefore, continuous analysis in randomized blocks with multi-year cropping would be important.

In conclusion, productivity tests of the Koshihikari homologous genotypes, the late *Hd16* and the short-statured, late *d60Hd16*, were conducted in nine prefectures in Japan and the relationship between genotype, traits and temperature accumulation was analyzed. *d60* and *H16* genotypes expressed stable traits adapted to a wide range of Japanese climatic conditions and growing environments. Furthermore, the results of various biostatistical analyses showed that quality was positively correlated with panicle length and 1000-grain weight, and yield was strongly positively correlated with 1000-grain weight. *d60* genotype was negatively correlated with culm length and lodging degree. Furthermore, the results of principal component analysis showed that in Koshihikari d60Hd16, Yamanashi and Ehime, which have longer panicle length and culm length, are better in terms of yield and quality, while Shimane, which is warmer and has shorter ear length and culm length, is better in terms of value of taste. The results also suggest that Koshihikari d60Hd16 can express traits that make it less susceptible to lodging degree while maintaining the same quality and yield as the wild type when grown as a late flowering line. This study will provide fundamental information for promoting smart agriculture using improved varieties.

## Acknowledgement

The authors express gratitude to the Startup Supporting Program (funding agency: Bio-oriented Technology Research Advancement Institution (BRAIN)) under Grant Number JPJ010717.

## Author Contribution

Conceptualization, M.T.; methodology, M.T., H.H.; data curation, M.T., H.H.; investigation M.T., H.H.; resources, M.T.; writing—original draft preparation, M.T., H.H.; writing—review and editing, M.T., H.H.; project administration, M.T.; funding acquisition, M.T.

## Supplementary Materials

**Supplementary table S1.**
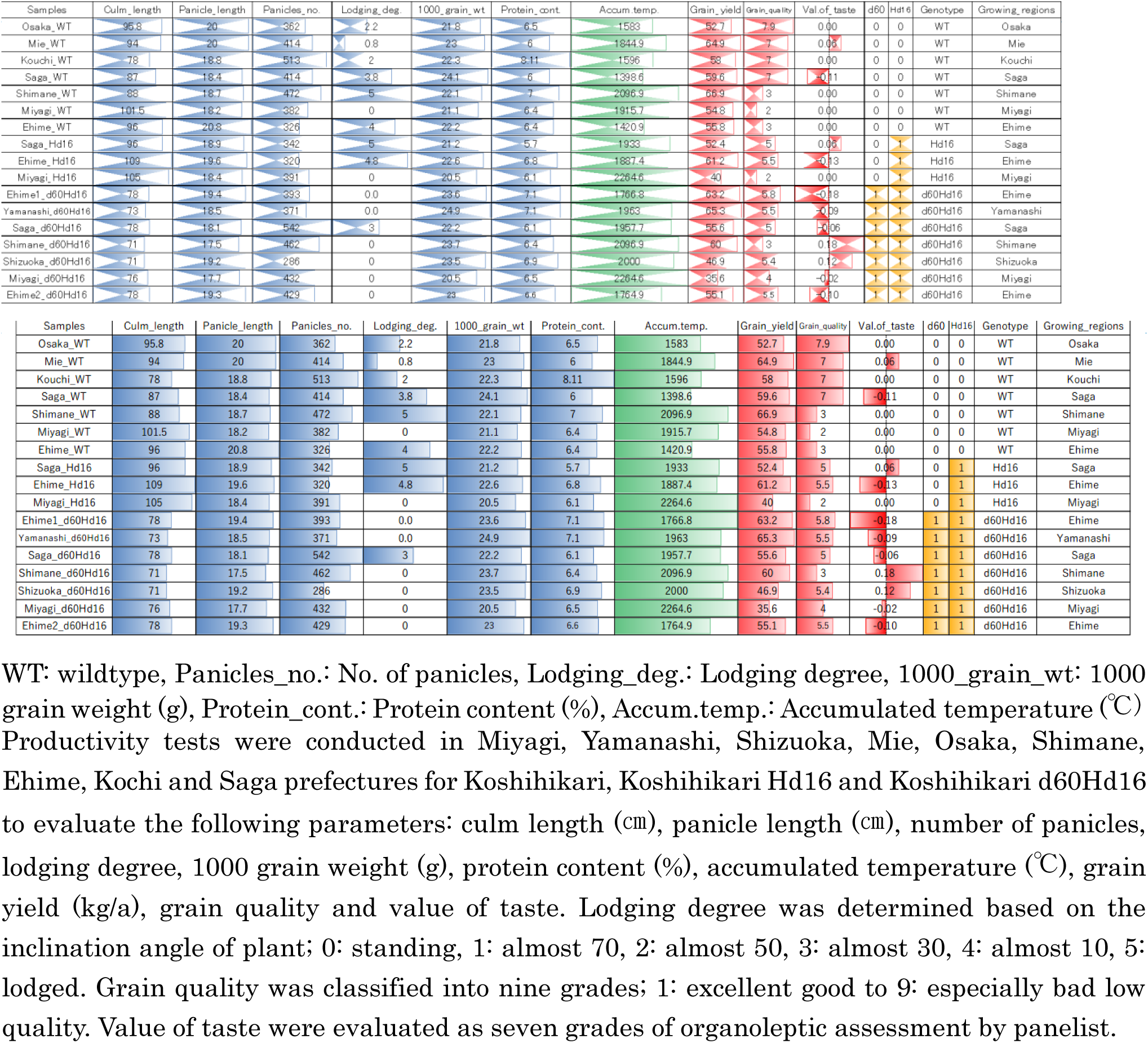
Summary of productivity test.

**Supplementary figure S2.**
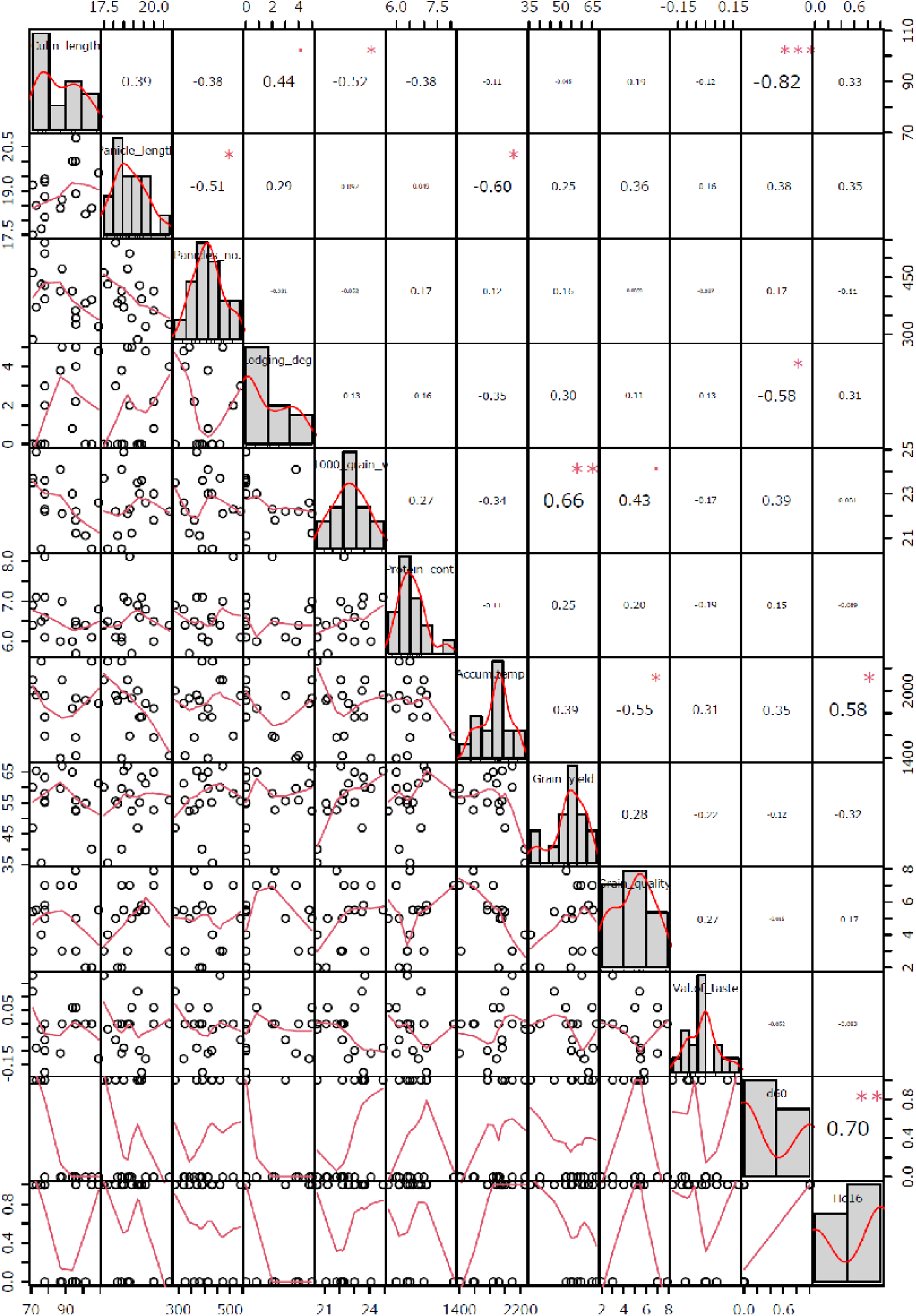
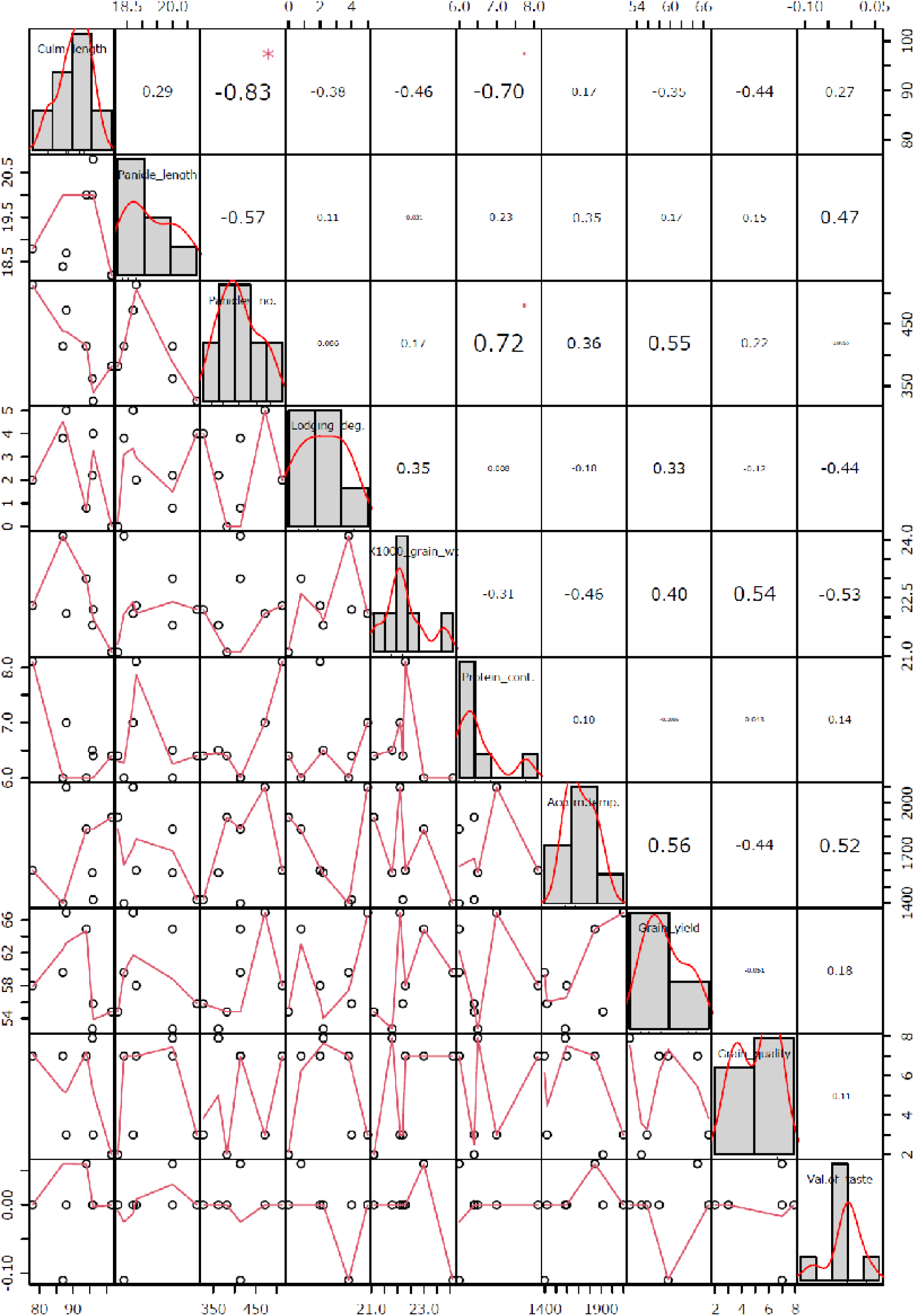
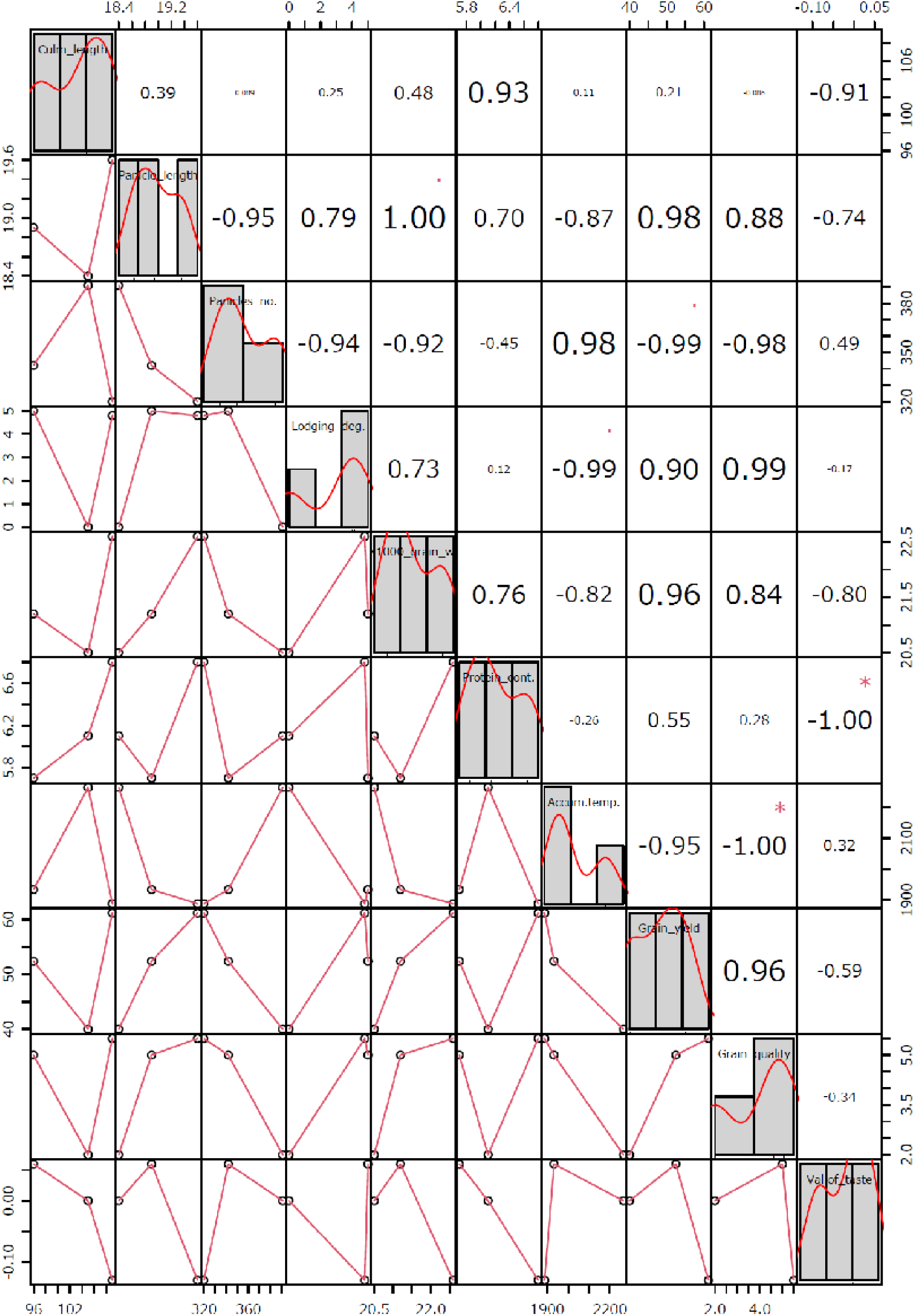
Scatter plots and correlation analysis. WT: wildtype, Panicles_no.: No. of panicles, Lodging_deg.: Lodging degree, 1000_grain_wt: 1000grainweight, Protein_cont.: Proteincontent, Accum.temp.: Accumulated temperature. A) all strains, B) wildtype, C) *Hd16*, D) *d60Hd16*.

**Supplementary figure S3.**
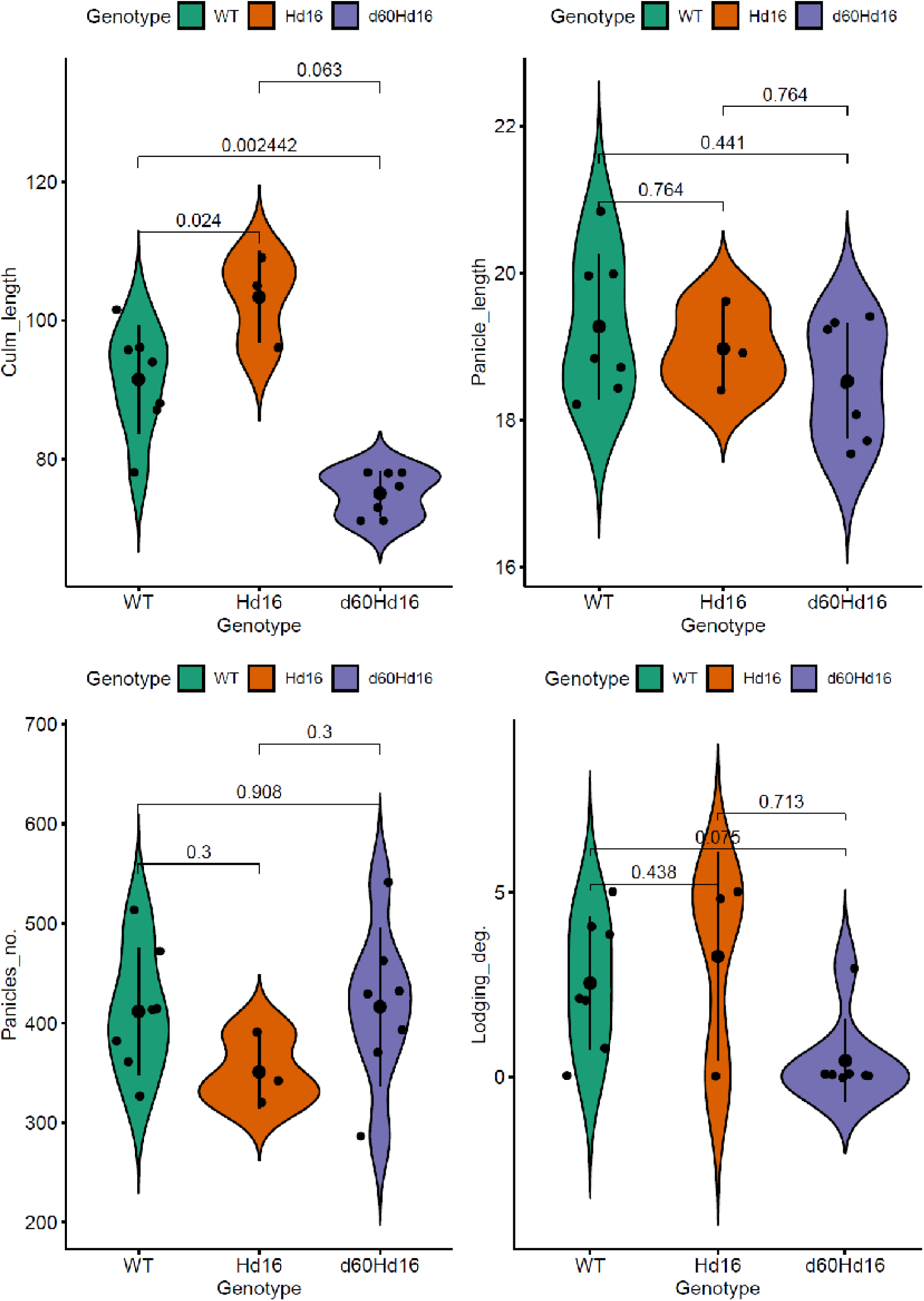
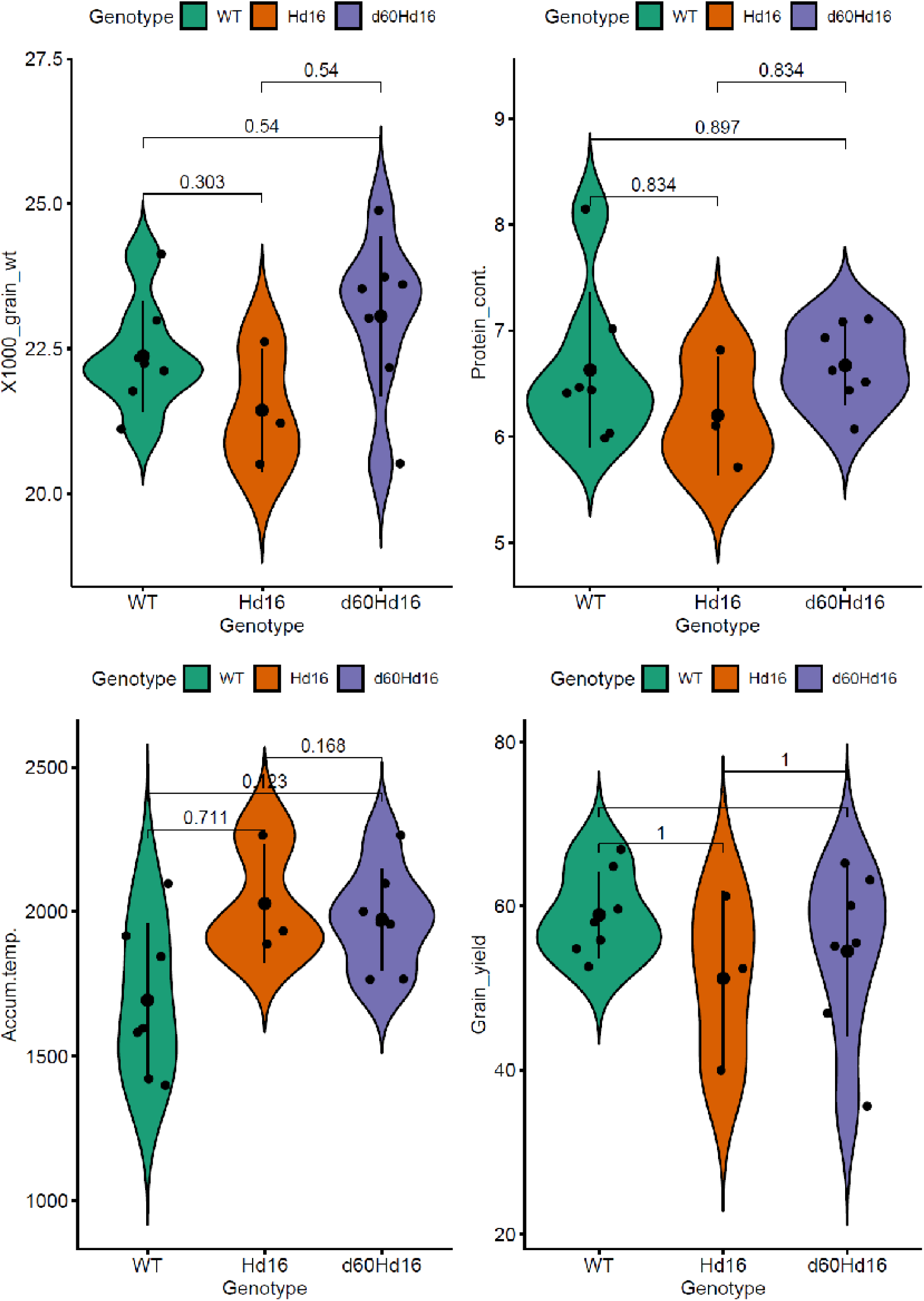
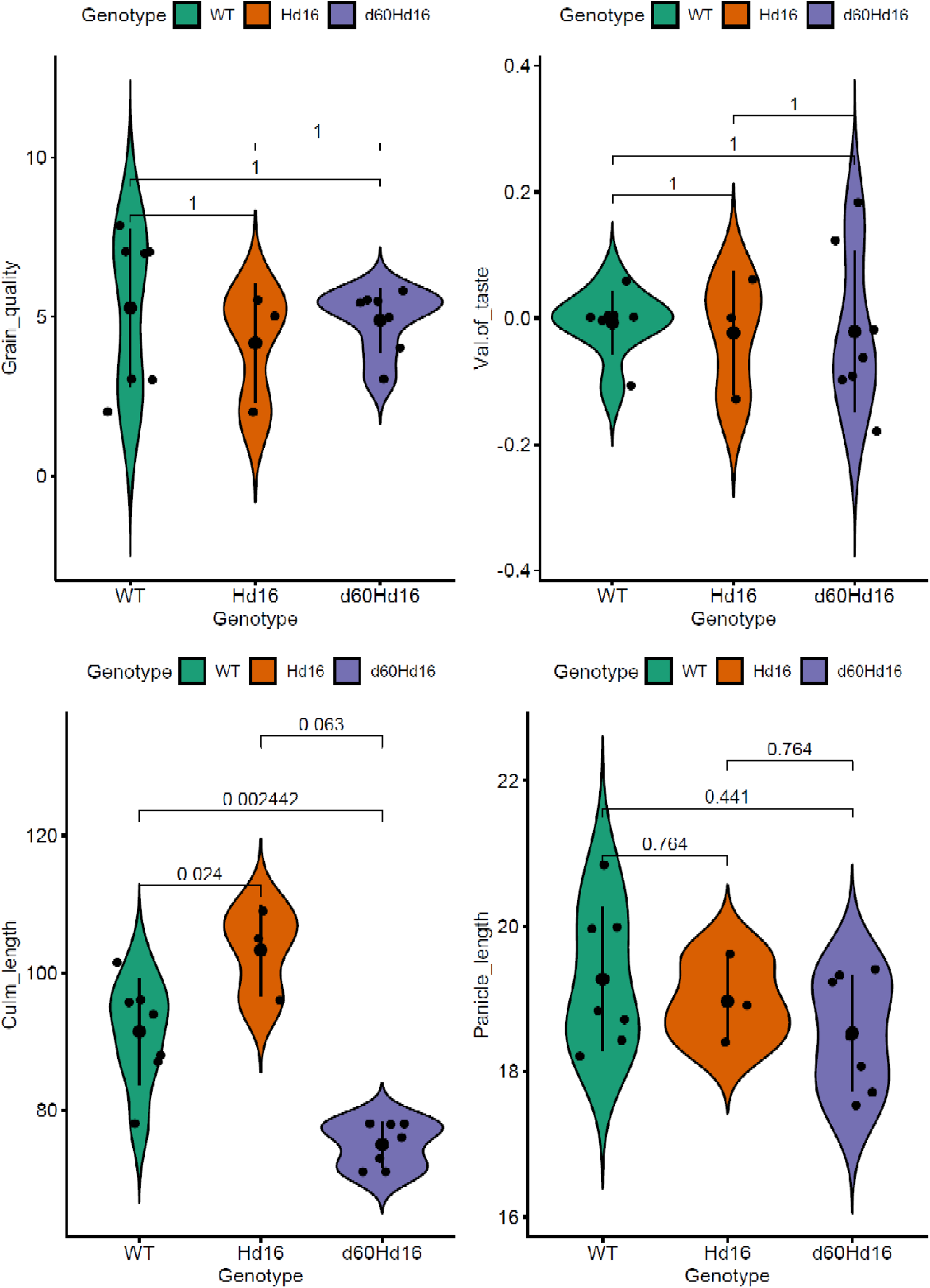
Comparison of traits, environmental factors, and target traits among genotype. WT: wildtype, Panicles_no.: No. of panicles, Lodging_deg.: Lodging degree, 1000_grain_wt: 1000_grain_weight, Protein_cont.: Protein content, Accum.temp.: Accumulated temperature

